# Alignment of Single-Molecule Sequencing Reads by Enhancing the Accuracy and Efficiency of Locality-Sensitive Hashing

**DOI:** 10.1101/2022.05.15.491980

**Authors:** Hassan Nikaein, Ali Sharifi-Zarchi

## Abstract

Aligning single-molecule sequencing (SMS) reads to a reference genome has been computationally challenging due to the high sequencing error rates in this technology. Short distances between consecutive errors in SMS reads confront finding seeds, subsequences of the reads with exact matches to the reference, that specifically target a unique genomic position. To overcome this issue, one can look for similarities, rather than exact matches. MinHash, a locality-sensitive hashing (LSH) scheme, measures the similarity of two sequences by listing all *k*-mers of each one and approximating the fraction of common *k*-mers between them using a family of hash functions, which usually includes hundreds to thousands of different hash functions in order to increase the measurement accuracy. MinHash is used to address various bioinformatics problems, including the assembly of SMS reads. Here, we enhance both the efficiency and accuracy of the MinHash scheme by algorithmic techniques. We use a single hash function, rather than hundreds or thousands of different hash functions as used in the other MinHash-based algorithms, without losing the accuracy. We also double the size of the seed sequences by allowing one sequencing error of any form inside a pair of *k*-mers, which has a significant impact on the accuracy. We show algorithm, called Aryana-LoR, outperforms the accuracy of the other existing SMS aligners in both E-coli and Human genomes.

**Availability:** Aryana-LoR is freely available at https://gitlab.com/hnikaein/aryana-LoR

## 1. Introduction

Single Molecule Sequencing (SMS) technologies created new hopes to reach fast and cost-effective amplification-free, massively parallel sequencing that can generate long reads that are critical for genome assembly and finding large structural variations [1, 2]. A major challenge to reach that ideal has been the high sequencing error rate, in forms of insertion, deletion, and substitution [3], due to an unavoidably low signal-to-noise ratio. Aligning SMS reads to a reference genome is particularly crucial for detecting structural variations, but the elevated sequencing error rate makes the problem computationally challenging [4, 5, 6].

In absence of large exact matches between SMS reads and the reference genome, one can look for non-exact matches through Locality-Sensitive Hashing (LSH) [7, 8]. For a given set of elements *S* with a metric distance function *d*, an LSH is a family *F* = {*h*_1_, *h*_2_, ···, *h_H_*} of *H* different hash functions. Each hash function *h_i_* maps the elements of *S* to a finite set of buckets *B*. Furthermore, there are two distance thresholds *δ*_1_ and *δ*_2_ (*δ*_2_ > *δ*_1_) and two probability values *p*_1_ and *p*_2_ (*p*_1_ > *p*_2_) such that for any hash function hj and any pair of elements *a, b* ∈ *S*:

- *d*(*a, b*) ≤ *δ*_1_ ⟹ *P*(*h_i_*(*a*) = *h_i_*(*b*)) ≥ *p*_1_, and
- *d*(*a, b*) ≥ *δ*_2_ ⟹ *P*(*h_i_*(*a*) = *h_i_*(*b*)) ≤ *p*_2_.

In other words, there is a high chance that *h_i_* maps the elements *a* and *b* to the same bucket if and only if they are highly similar. LSH functions have received attention in different areas of data science. One of their first applications in bioinformatics was to perform an ungapped comparison of long sequences [9]. MHAP algorithm employs LSH for assembly of the SMS reads [10]. Briefly, MHAP uses a widely-used LSH scheme called MinHash [11]. Given two sequences *A* and *B*, MinHash rolls a window of size *k*-bp over each one to create *shingles*, overlapped *k*-mers. Then it approximates the Jaccard similarity of *A* and *B* by a family of *H* different hash functions, as explained in the Methods section. MinHash scheme is also recently used in Mashmap2, an algorithm for computing whole-genome homology maps [12]. An earlier version of it, called Mashmap, was used to approximate the alignment location of SMS reads to a reference genome [13].

Here we employ LSH for the alignment of SMS reads to a reference genome, using two techniques that enhance both accuracy and efficiency of MinHash scheme for long and noisy reads. The enhanced accuracy is obtained by using a pair of shingles by allowing a sequencing error, in all forms of insertion, deletion, and substitution, inside them. Furthermore, the efficiency of the MinHash sketching is improved by a factor of *H* (which is 512 to 1 256 for different configurations of MHAP and 200 to 500 for Mashmap) by using *one permutation hashing* technique [14].

## 2. Methods

### 2.1. MinHash

The central aim of MinHash is to approximate the Jaccard similarity coefficient *J*(*A, B*) between two sets *A* and *B* (*A, B ⊂ S*): 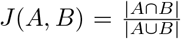. The Jaccard similarities are 0 and 1 for two pairs of disjoint and equal sets, respectively. Otherwise, the value is strictly between 0 and 1 and two sets are more similar if their Jaccard similarity is closer to 1. If *h* is a collision-free hash function, that maps distinct elements of *S* to different discrete values, and for a given set *X* ⊂ *S* we define *h*(*X*) = {*h*(*x*)|*x* ∈ *X*} and *h*_min_(*X*) as the minimum value of *h*(*X*), then *h*_min_(*A*) = *h*_min_(*B*) iff the element having the minimum hash value in *A* ∪ *B* is present in both *A* and *B*. In other words:

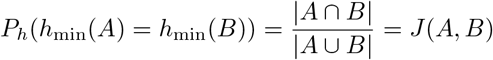

Let *I* be an indicator random variable that has a value of 1 if *h*_min_(*A*) = *h*_min_(*B*), or 0 otherwise. The above relation shows that the expected value of *I* is *J*(*A,B*). Hence, *I* is an unbiased estimator of the Jaccard similarity. It has, however, a high variance since the values of *I* are only 0 or 1.

To address this issue, MHAP uses a family of *H* different random hash functions. For each sequence *A*, it generates a *sketch*, which is (min_1_(*A*), min_2_(*A*), ···, min_*H*_(*A*)). Here, min_*j*_(*A*) is a simpler notation for 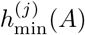 where *h_(j)_* is the *j*-th hash function. The value of *J*(*A,B*) is then approximated by:

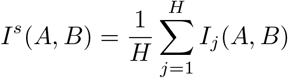

where *I_j_*(*A, B*) is the indicator variable having a value of 1 if min_*j*_(*A*) = min_*j*_(*B*), or 0 otherwise; And *I^s^* is an unbiased estimator of the Jaccard Similarity with smaller variance. It is trivial to show that expected value of *I^s^*(*A, B*) is *J*(*A, B*) and 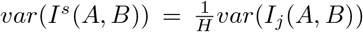. Therefore Minhash method with *H* function is *H* times more accurate than j with one function.

The time complexity of this MinHash scheme is *O*(*LH*), where *L* is the maximum length of the given pair of sequences. Here the size of shingles (*k*) is considered to be much smaller than *L*.

### 2.2. Gapped shingles

There are two challenges in using shingles of MinHash for SMS reads: (I) If the size of the shingle is too short, it will have a lot of exact matches in the reference genome; hence, the outcome might not be accurate enough. (II) By increasing the size of the shingle, there will have a higher chance that hits by sequencing errors.

If we consider a uniform and independent sequencing error of *p_e_* per nucleotide, a shingle of length *k* will be hit by sequencing error with a probability of 1 – (1 – *p_e_*)^*k*^. On the other hand, if we consider the reference genome as a random sequence of length *G* with independent and uniform distribution of bases, the expected number of random occurrences of a shingle of size *k* in the reference genome will be 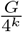. As a result, a *k* = 7 bp shingle has a 32% chance of surviving a 15% SMS error rate per nucleotide (the sum of mismatch, insert and delete error rates); but it will have an expected number of 183 105 exact matches in the human reference genome. On the other hand, a *k* = 14 bp shingle will have about 11 hits in the human reference genome; but it is 90% likely to be hit by a sequencing error. Hence, neither of shingle lengths work great.

To address this issue, we extended the idea of shingles in MinHash to gapped shingles, which for simplicity we call them here as *gingles*. Each gingle is a sequence of two or more shingles with a specific number of gaps between them. In this paper, we consider a *k* bp gingle as two *k*/2 bp shingles that are one bp apart. For instance, if a fragment of DNA is `“ACGTGGA”`, a 6 bp gingle would be `“ACGGGA”`, in which the nucleotide `“T”` between the two 3 bp shingles is removed.

To have both benefits of both short shingles, which are reasonably error-free, and large shingles that are significantly more specific in the reference genome, we can use a combination of shingles and gingles. Consider a particular region of a read, and the corresponding part in the reference, and assume there will be at most one sequencing error of size 1 bp. We create *k* = 6 shingles and gingles of both the reference and the read. If the read matches the reference, the shingles of both sequences will match. If the read has an insertion, a gingle of the read that has the insertion in its gap will match a reference shingle. One bp deletion in the read will be handled by a match between a reference gingle and a read shingle, and a mismatch is handled by matching the gingles of both sequences.

As a proof-of-concept, we examined whether this idea is effective by simulating 10 000 bp reads, with one sequencing error (insertion, deletion or mismatch) after each correctly sequenced interval. The sizes of such intervals were chosen between 6 to 8 bp randomly. This error pattern is very challenging for seedbased or shingle-based alignment methods since the seed sizes or shingles are very limited. As expected, a min-hashing algorithm that used only shingles could not accurately identify the genomic location of each read, due to the very small size of correctly sequenced intervals. On the other hand, a combination of 12 bp gingles and shingles could determine, at a reasonable accuracy, the correct genomic location of the simulated reads.

### 2.3. Fast MinHash using baskets

The running time of the classic MinHash is linearly increased by increasing *H*, the number of different hash functions. The lower number of hash functions results in improved efficiency, at the cost of an increased variance of the whole set of indicator variables, and the reduced accuracy of the approximated Jaccard similarity.

We used a basket-min idea which has been introduced first time in [14] with the name “one permutation hashing”. This idea uses a single hash function *h* instead of *H* different hash functions, but divides its domain into *H* parts to obtain the same accuracy with *H* hash functions version. Suppose *h* is (almost) uniformly distributed over a range of [0, *R*]. We define *n* baskets *b*_1_, *b*_2_,···, *b_n_* where 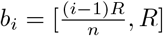. Given a sequence *A* with a set of gingles (or shingles) *G*, a sketch of size *n* is defined as 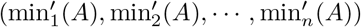 where: 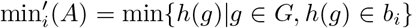. We define 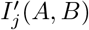 as the indicator variable with a value of 1 in case of 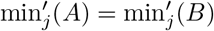 and 0 otherwise. As a result we define:

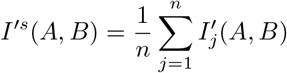

We will show that this method accuracy is the same as the original Minhash with *n* hash functions. (Also, [14] showed this with a different proof.)

#### Lemma 1.

*The expected value of I’^S^(A,B) is equal to J(A,B).*

Proof. Since *h* is (almost) uniformly distributed over 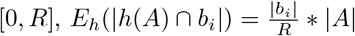. Therefore

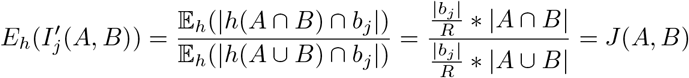

Therefore

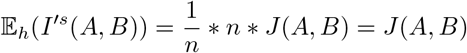

#### Lemma 2.

*The variance of I’^S^(A,B) is equal to 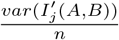.*

Proof. First we show that 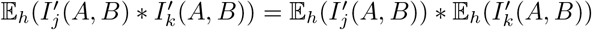.

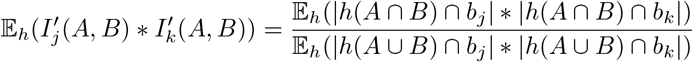

Since *h* is (almost) uniformly distributed over [0, *R*], |*h*(*A* ⋂ *B*) ⋂ *b_j_*| is independent from |*h*(*A* ⋂ *B*) ⋂*b_k_*| and |*h*(*A* ∪ *B*) ⋂ *b_j_* is independent from |*h*(*A* ∪ *B*) ⋂ *b_k_*|. Therefore:

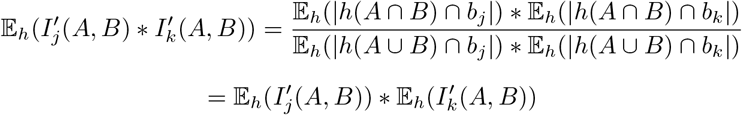

So, we have:

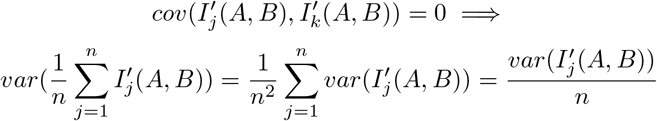

### 2.4. Tag selection

The overall SMS read alignment algorithm breaks the reference genome into overlapping tags of various sizes. For simplicity of calculations, we select tag sizes as powers of two. For a tag size *T*, the reference genome is covered with two sets of consecutive tags, one set starting from the very first nucleotide of the genome and the other set starts at nucleotide *T*/2 + 1-th (Fig. 1A). This is to ensure at least one tag will cover the whole genomic region of each read. The set of selected tag sizes is {2^9^, 2^11^, 2^13^, 2^15^} based on the distribution of SMS reads, obtained from real Pacific Biosciences and Oxford Nanopore datasets. Each tag is assigned a unique label. To increase Jaccard similarity between the read and the tag, we compare each read against the smallest tags that have a size more than or equal to the read.

**Figure 1:**
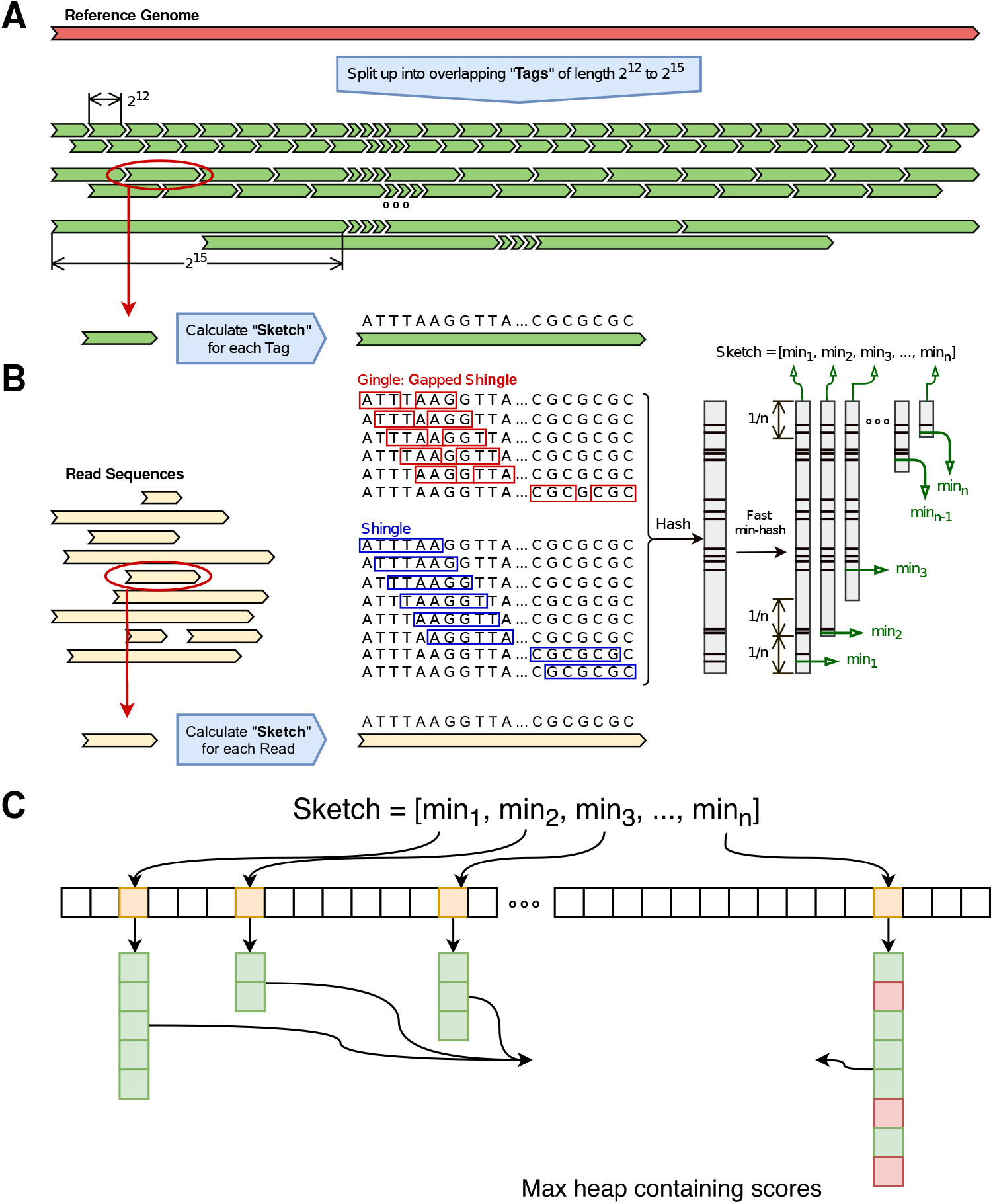
Algorithmic scheme of finding the genomic position of SMS reads. (A) The reference genome is split into overlapped tags of different lengths. The min-hash sketch of each reference tag is computed. Each tag has a unique label. (B) The min-hash sketch of each read is also computed. All shingles and gingles are hashed. The whole range of hash values is split into different baskets. The min value of basket *k* is the *k*-th element of the sketch. (C) For each value of the sketch, the reference index holds the list of labels of all tags which have the same min value for the same sketch basket. If the length of such list is too long, a smaller set of random tag labels are kept (those in green). For each read, the tags having the maximum score are stored in a max-heap priority queue.

We compute the sketch of all gingles and shingles of every tag. We do the same for every read (Fig. 1B). The sketches are computed based on our baskets scheme. We used a reversible hash function, similar to Minimap, which is proven to have zero collisions. This had a significant effect on the accuracy of the algorithm. The code of the hash function is available in Aryana-LoR source code.

The index file, for the reference genome, is created during a pre-process (Fig. 1C). We create an array of size *R* + 1, if all values of the hash function fit in the range of [0, *R*]. Each cell *i* of the index array holds a pointer to a list of tag labels that have the value *i* somewhere in their sketch. If due to the overlaps among the baskets, the value *i* appears multiple times in the sketch of a tag, the tag label is added only once to the list pointed by the cell *i*. If the list of cell *i* is too long (e.g., for repeat regions), only a subset of tag labels can be kept depending on the parameters of creating reference index.

For aligning each read, we create an empty max-heap priority queue. This queue holds the tags that have some sketch values in common with the reads. The tag having the maximum number of matched sketch values is always kept and dynamically updated in the root of the heap. For each position *j* of the read sketch (1 ≤ *j* ≤ *n*), we go to the cell *min_j_* of the index array and retrieve the list of tags having the same value in their sketches. We increment the score of each tag in the list and update its position in the priority queue. Finally, a subset of tags with the highest scores is selected as candidates. We then select the best candidate using Aryana-LoR, as follows.

### 2.5. Aryana

The algorithmic approach of Aryana for aligning short reads is already published [15]. Briefly, it splits the reference genome into equal size intervals called *tags*. For each read, it tries to locate all maximal length seeds (read substrings that exactly match to the reference genome) using Borrows-Wheeler Transform [16], and score corresponding tags based on the length of each seed. A hash table keeps the total scores of each tag for a read, along with the read number to eliminate the need to reset the hash table for each new read. Several tags with the maximum total score are selected as candidates, and for each candidate, a maximal chain of seeds between the read and the tag is identified. A modified Smith-Waterman algorithm computes the alignment penalty for each pair of gaps, one in the reference and another in the read, between each two consecutive seed in the maximal chain. Finally, it reports the least penalized alignment (or a user-defined number of best alignments). More details are provided in the previous article [15].

For the SMS reads, if a tag *i* is selected as the candidate for a subset of the reads, Aryana-LoR takes tag *i* as the reference and aligns each read of the subset to it. Since the read might not completely fit the candidate tag, we double the size of the tag while treating it as the reference by extending it from both ends. Since the size of each tag is substantially smaller than the size of the whole reference genome, we can run Aryana-LoR with a very small minimum seed size. Although smaller seed sizes are not specific for the whole reference genome, they are distinct within a single tag. Hence the alignment of an SMS read to a tag can be performed much more accurately and efficiently than aligning it to the whole reference genome. We finally report, for each read, the best hit among all different candidate tags.

## 3. Results and Discussion

### 3.1. MinHash with the baskets scheme accurately approximates Jaccard similarity, with enhanced efficiency

To check how the baskets scheme works in practice, we simulated reads of different lengths (1000 and 10000 bp) and treated them with a Poissonian random noise of different rates (1% and 5%). Each dataset of reads contained 1000 reads. We developed maximally similar implementations of a classical MinHash algorithm consisting of *H* = 100 different hash functions, and a basket MinHash algorithm with *H* = 1 hash function and *n* = 100 baskets. By measuring the actual Jaccard similarity between each read and the related genomic position, we compared the similarities approximated by each method.

As shown in Fig. 2, the basket scheme MinHash outperforms the classical MinHash when the shingle size is shorter than 9 or 10 bp, depending on the length of the reads and the error rate. Average of absolute difference of basket scheme MinHash and real Jaccard similarity is better than regular min hash because the latter overfits and therefore have more variance than basket scheme MinHash. For longer shingles, the accuracies are comparable, particularly for the 10000 bp read. Due to the high current error rate of SMS reads (15%), we know the average size of shingles should be lower than 9 to keep a reasonable part of the shingles error-free. This shows that reducing the number of hash functions to 1 in basket scheme does not affect the accuracy of the aligner.

**Figure 2:**
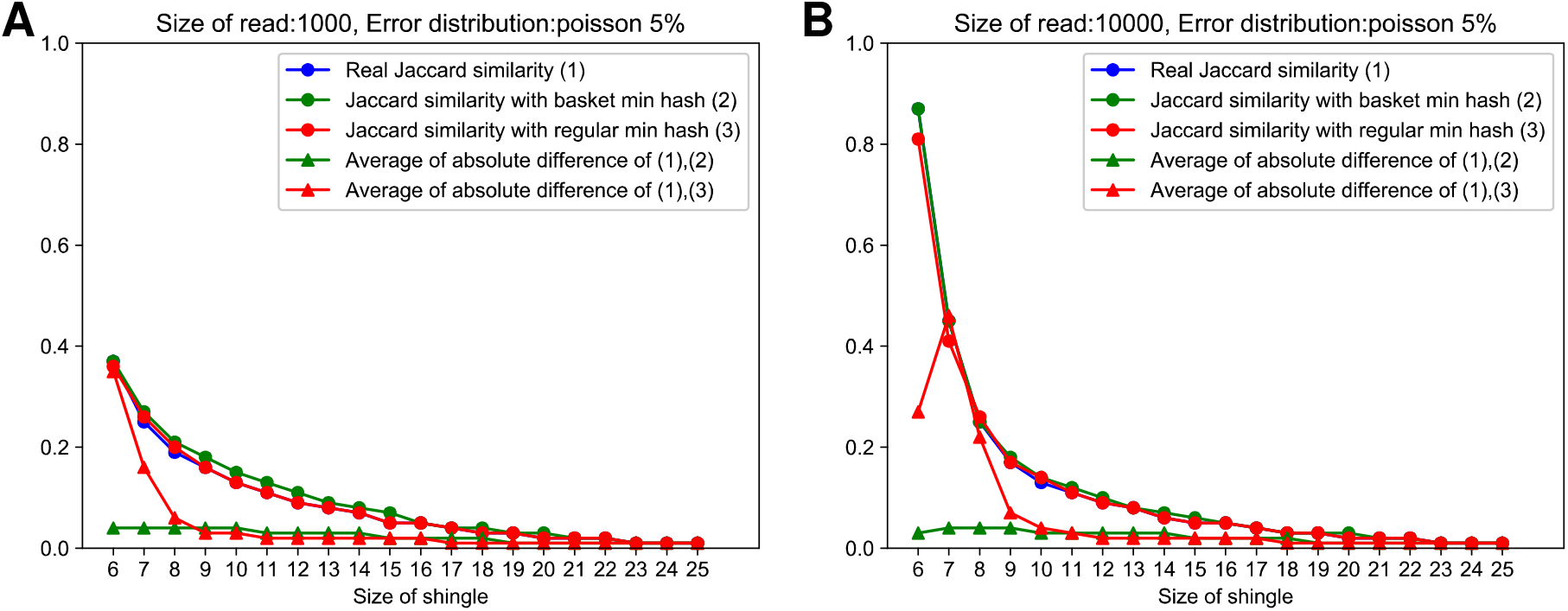
Basket scheme MinHash accurately measures Jaccard similarity. Simulated 1000 bp (A) and 10000 bp (B) reads are treated with Poisson errors with 5% error rate. The line plots with rounded dots show the correct Jaccard similarity between each read and the representing region in the genome (blue) and approximated values by using basket scheme with a single hash function (green) and classical MinHash using 100 different hash functions (red). The line plots with triangles show the average absolute error of the basket scheme (green) and the classical MinHash (red).

Furthermore, by using only a single hash function the speed of the sketching part of the aligner would be increased by a factor of *H*. For instance, MHAP uses *H* = 512 for the fast and *H* = 1256 for sensitive configurations. Hence using the basket scheme can improve the sketching time, which is one of the bottlenecks of the alignment running time, by three orders of magnitude.

### 3.2. Simulating reads based on real SMS data

To compare the accuracy of Aryana-LoR with other SMS read aligners, we selected three other SMS aligners: (I) Mashmap, (II) Minialign, a PacBio and Nanopore aligner based on the seed-and-extend strategy, minimizer-based index of the Minimap overlapper, array-based seed chaining, and SIMD-parallel Smith-Waterman-Gotoh extension [17], and (III) Minimap2, a program that aligns DNA or mRNA long-read sequences that uses a hash table of minimizers, sorts and chains them with dynamic programming and finds global alignment between internal seeds [18].

To measure the accuracy of each method, we required to simulate reads that were very similar, in terms of different statistical parameters, to the actual reads. For this purpose, we obtained Pacific Biosciences (PacBio) Human54x datasets, which is a 54x coverage human SMS data of *de novo* human sequence assembly from the website of the company [19]. The NA12878 data of Oxford Nanopore Whole-Genome Sequencing (WGS) consortium was obtained from GitHub [20].

By testing a number of SMS read simulators, we identified SimLoRD [21] and PBSIM [22] to have better performances. By generating the distribution of eight different statistical parameters for the real and the simulated data, we observed simulators do not match the real data in two parameters: the distributions of read lengths, and also the longest size of seed in each read (Fig. 3).

**Figure 3:**
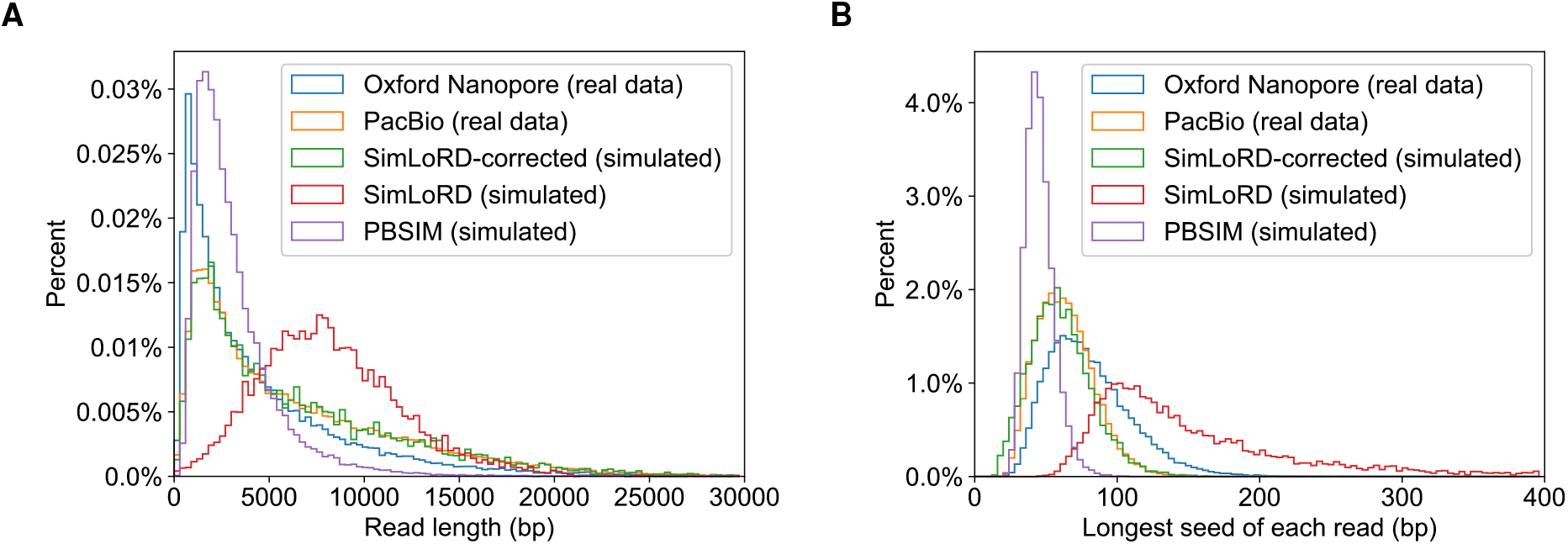
Simulating reads based on the real SMS data. Two sets of histograms are provided, depicting the distributions of read lengths (A) and the longest seed of each read (B) in real and simulated datasets.

To address this issue, we provided SimLoRD a file, as an input argument, containing reads sizes that matched the real data. We also changed the source code of SimLoRD to adjust the length of the seeds (error-free subsequences of the read) according to a Gamma-distribution that was fitted to the real data. The parameters of the Gamma distribution are adjusted to the real data. As depicted in Fig. 3, the SimLoRD-corrected has the most similar distributions to the real PacBio and Nanopore data.

### 3.3. Comparing SMS aligners on simulated data

Using 10 000 simulated reads from both E-coli and human genomes, we compared the alignment results of Mashmap, Minimap2, Minialign, and Aryana-LoR. In addition to the normal configuration of Aryana-LoR that used both shingles and gingles, we also tested a particular configuration that used only shingles.

It is very important to mention that for each aligner, we carefully set all of available configurations and arguments such that the maximum accuracy is obtained. This permitted us to compare the maximum accuracy of all aligners.

As shown in Fig. 4, Aryana-LoR outperformed the other aligners based on the accuracy of the alignment position. For each simulated read, we computed the absolute genomic distance between the correct location of the read and the alignment position reported by each SMS aligner. Then we used different thresholds as the maximum absolute distance to consider the read is correctly aligned. These thresholds included 10, 100 and 1000 bp, and also half of the read size. Although the fourth threshold is too large and variable, we included it in our analyses because: (I) Mashmap used that parameters for computing its accuracy, (II) this threshold ensures the read has partial overlap with the alignment region.

**Figure 4:**
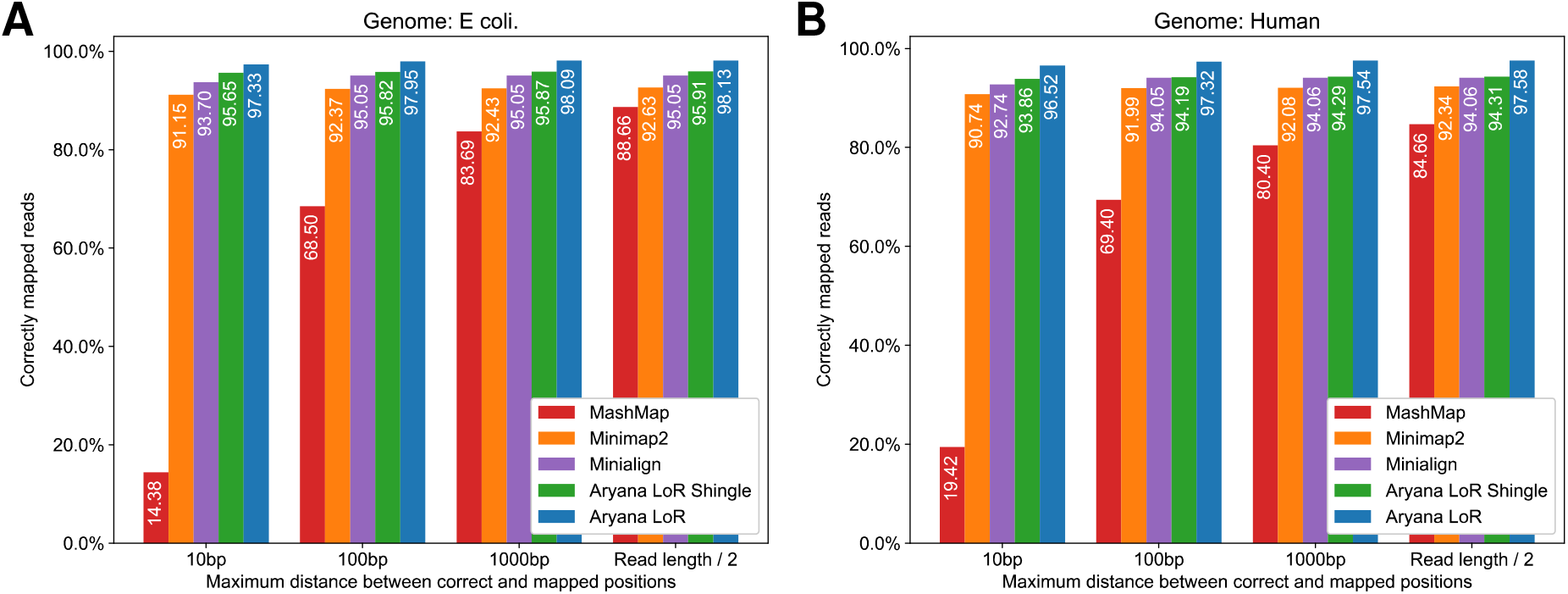
Comparing the accuracy of the alignment position for simulated reads from E-coli (A) and human (B) genomes using different SMS aligners. In each plot, the x-axis shows different maximum thresholds for the absolute distances between the alignment position of each read from its original position in the genome. The y-axis shows percent of aligned reads having an absolute distance lower than each threshold value. Different aligners are color-coded.

Fig. 4A show the results for E-coli genome using default penalty arguments of each aligner. As shown, in all comparisons both Aryana-LoR and Aryana-LoR with shingles outperform other aligners in all threshold values for maximum absolute distance. Minialign, Minimap2, and Mashmap have the third, fourth and fifth rankings in all comparisons, respectively. For the smaller absolute distance thresholds such as 10 bp and 100 bp, Mashmap has a significant accuracy distance with the other aligners, but this gap is reduced when the half of read length is taken as the maximum distance between aligned and correct positions.

Another important observation is the elevated accuracy of Aryana-LoR by using both shingles and gingles. It confirms that the gapped shingles method has a significant impact on the accuracy of the aligner while using MinHash technique. The elevated accuracies are between 1.66% to 2.22%, based on different distance thresholds.

Fig. 4B show similar analyses for the simulated reads from the human genome. Again, both Aryana-LoR and Aryana-LoR with shingles have the best performances in all comparisons. The accuracy of Minialign is very close to Aryana-LoR with shingles for 100 and 1000 bp thresholds when each aligner is using its default penalty arguments, but still, there is a significant accuracy gap between Aryana-LoR using both gingles and shingles with the other aligners. The accuracy distance between Aryana-LoR and Aryana-LoR-shingles is also increased from 2.66% to 3.27%, depending on the distance threshold.

### 3.4. Comparison of the aligners on real data

We compared Aryana-LoR with Minialign and Minimap2 using 1000 real Oxford Nanopore reads, obtained from NA12878 dataset of the Whole Genome Sequencing Consortium [20] and 1000 Pacific Biosciences reads obtained from Human54x datasets from the website of the company [19].

Since the correct positions for the real reads are unavailable, our criterion was to measure the total match score produced by each algorithm. Mashmap only reports the alignment position without the CIGAR sequence, hence it was excluded in this analysis. Fig. 5A, B show the pairwise comparison of Aryana-LoR with Minimap2 in the human genome for PacBio and Nanopore reads. Fig. 5C, D show the same comparisons between Aryana-LoR and Minialign. In each plot, the reads are depicted by dots, and their sizes are shown by the color (the longer reads are darker). To enable the comparison of all aligners, we had to assign similar parameters for penalties to all aligners. We used (1, 3, 3, 2) as the match score, mismatch penalty, gap open and gap extension penalties, respectively, as the input parameters of all aligners. Furthermore, the total match score of each read was computed using the same coefficients. Normalized scores were computed by dividing the match score of each read by the length of reads. We set the minimum match score of −1.2 to all unaligned reads by each aligner, for visualization purpose.

**Figure 5:**
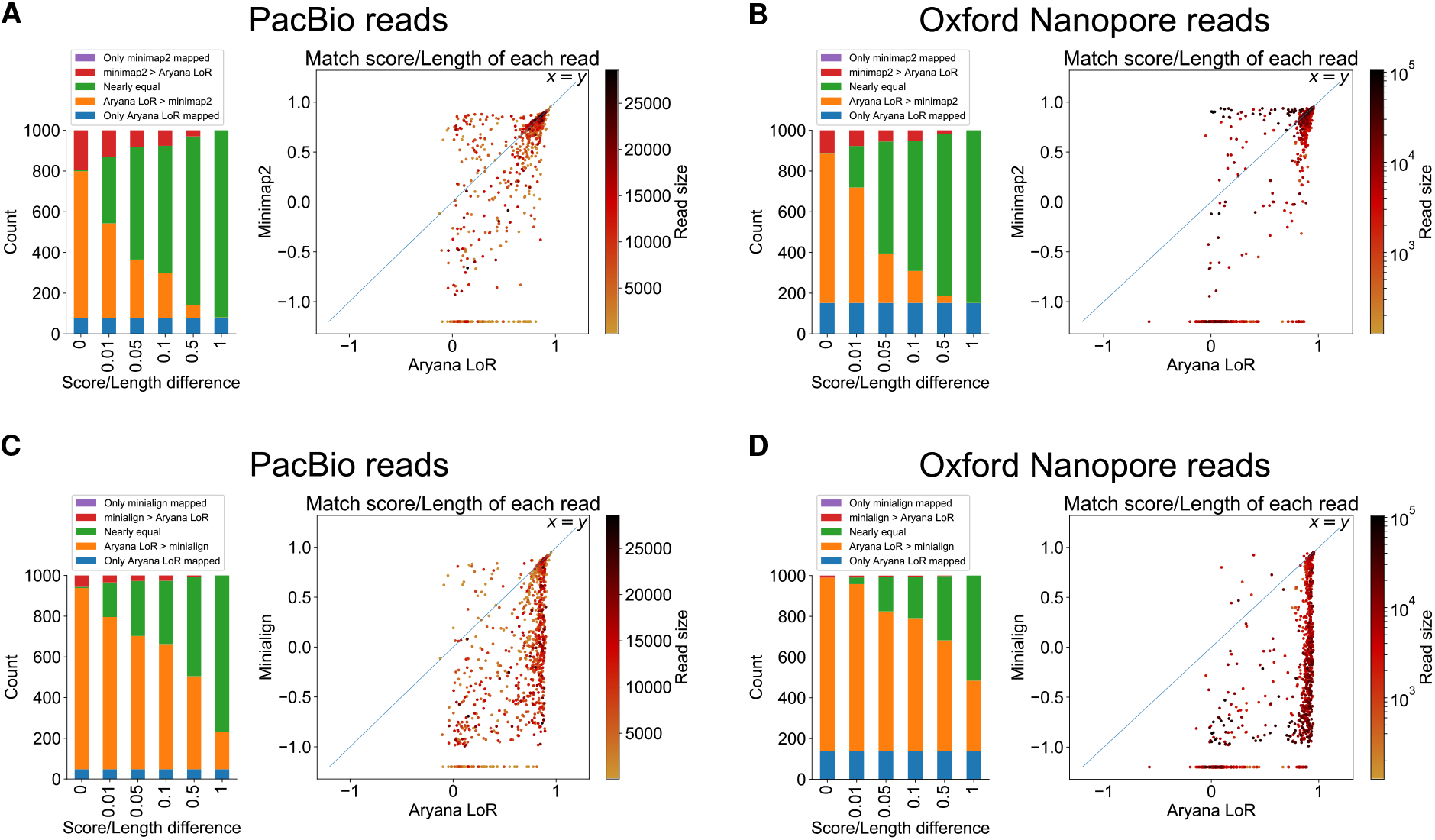
Performance of Aryana-LoR in comparison with other aligners on 1000 real human SMS reads. For each read, normalized alignment score is computed by dividing the alignemnt score of each algoirthm by the read length. For barplots, the x-axis shows different thresholds for comparison of the normalized scores. For each threshold *t*, green color represents reads that their normalized scores between Aryana-LoR and the other algorithm have at most *t* units distance. Orange and red show reads that have normalized score higher for Aryana-LoR or the other aligner, respectively. Blue and purple show reads that are only aligned by Aryana-LoR or the other aligner, respectively. In scatter plots, each point represents one real read; the axes show the normalized scores using Aryana-LoR (x-axis) and either Minialign or Minimap2 (y-axis). The color of each point shows the length of the read. (A, B) Comparison of Aryana-LoR versus Minimap2, (C, D) Aryana-LoR versus Minialign. (A, C) using real Pacific Biosciences(PacBio) reads. (B, D) using real Oxford Nanopore reads.

As shown in Fig. 5A, B, Aryana-LoR has superior results than Minimap2 in both PacBio and Oxford Nanopore reads. Fig. 5C, D shows Aryana-LoR outperforms Minialign. Comparing the results of Minimap2 and Minialign shows that Minialign gives a better position for simulated reads, however, in real reads Minimap2 have better performance.

### 3.5. Comparison of the running time and memory usage of different aligners

Tables 1 and 2 show the memory and time usages of different aligners while analyzing 10 000 simulated reads of the E-coli and human genomes, respectively. For shorter genomes, Aryana-LoR consumes more memory due to the structure of index files. For larger genomes such as human, however, it uses the smallest amount of memory, which is 2GB-25GB less than what is used by the other aligners. The memory usage of Aryana-LoR is mainly due to index files, hence increasing the number of reads does not affect it significantly.

**Table 1:** Time-Memory suite

| E coli. | Aryana LoR | MashMap | Minimap2 | Minialign |
| --- | --- | --- | --- | --- |
| Memory (MB) | 2142 | 75.76 | 229.74 | 87.71 |
| Time (s) | 329.88 | 20.09 | 25.81 | 5.58 |

**Table 2:** Time-Memory suite

| Grch38 | Aryana LoR | MashMap | Minimap2 | Minialign |
| --- | --- | --- | --- | --- |
| Memory (GB) | 13.42 | 38.894 | 19.618 | 15.511 |
| Time (s) | 5592 | 5549.1 | 250.26 | 151.82 |

The time usage of Aryana-LoR is considerably higher than the other aligners. For larger genomes such as human, there is a big gap between Aryana-LoR and Mashmap that both use the MinHash scheme on one side, and Minimap2 and Minialign which use minimizers on the other side. The higher speed of the latter algorithms comes at the cost of reduced and bounded accuracy. The MinHash scheme allows increasing the accuracy by increasing the number of hash functions, or as we suggested here, by using multiple baskets. Although Aryana-LoR and Mashmap have a similar running time on human reads, about half of Aryana-LoR time is consumed to generate the CIGAR sequences that contain the precise alignment, which are not produced by Mashmap. Hence we conclude that Aryana-LoR has a better performance than Mashmap for the longer genomes.

## 4. Acknowledgements

We are grateful for the support of Dr. Hamidreza Chitsaz (Colorado State University) and Dr. Rasool Jalili (Sharif University of Technology). Authors thank Amir Ali Moeinfar for the artworks of Figs. 1, Mohammad Roghani and Milad Aghajohari for their contributions in developing Aryana. Part of the analysis was performed on the computing cluster of the computer science department, Colorado State University.

